# Characterization of a Biochemical Mouse Model of Primary Aldosteronism for Thermal Therapies

**DOI:** 10.1101/2024.05.07.592955

**Authors:** Sarah A. Timmerman, Nathan Mullen, Angela E Taylor, Lorna C Gilligan, Marla Pyle, Tej B. Shrestha, Jan Sebek, Margaret A. Highland, Ritihaas Challapalli, Wiebke Arlt, Stefan H. Bossmann, Michael Conall Dennedy, Punit Prakash, Matthew T. Basel

## Abstract

**Introduction:** Aldosterone-producing adenoma (APA) is the most common cause of endocrine-related hypertension but surgery is not always feasible. Current medical interventions are associated with significant side effects and poor patient compliance. New APA animal models that replicate basic characteristics of APA and give physical and biochemical feedback are needed to test new non-surgical treatment methods, such as image-guided thermal ablation,

**Methods:** A model of APA was developed in nude mice using HAC15 cells, a human adrenal carcinoma cell line. Tumor growth, aldosterone production, and sensitivity to angiotensin II were characterized in the model. The utility of the model was validated via treatment with microwave ablation and characterization of the resulting physical and biochemical changes in the tumor.

**Results:** The APA model showed rapid and relatively homogeneous growth. The tumors produced aldosterone and steroid precursors in response to angiotensin II challenge, and plasma aldosterone levels were significantly higher in tumor bearing mice two hours after challenge verses non-tumor bearing mice. The model was useful for testing microwave ablation therapy, reducing aldosterone production by 80% in treated mice.

**Conclusion:** The HAC15 model is a useful tumor model to study and develop localized treatment methods for APA.

## 1. Introduction

Aldosterone-producing adenoma (APA) is a noncancerous adrenal tumor also known as Conn’s adenoma. In APA patients, somatic gene mutations generate hypersecretion of aldosterone independent of its primary regulators: angiotensin II and hyperkalemia. ^1–3^ PA (primary aldosteronism) manifests clinically as hypertension due to hypernatremia and hypokalemia. Of patients referred for hypertension, 6-13% have PA with 30-40% of those diagnosed with APA. ^4^ In addition, PA is now considered the most common cause of endocrine-related hypertension. Cardiac complications, including left ventricular hypertrophy, atrial fibrillation, and cardiac fibrosis, are common sequelae to PA hypertension and occur with greater frequency than in BP-matched primary hypertension. ^5^ Metanalysis has demonstrated that PA patients have a higher risk of developing other related conditions such as atrial fibrillation, stroke, and heart failure. ^6–9^ Furthermore, co-secretion of glucocorticoids is sometimes observed in PA (REF), ^10^ which is associated with significant co-morbidity including type 2 diabetes and other metabolic conditions (REFS). ^11–15^ Chronic high levels of aldosterone can impact mental well-being. Mineralocorticoid receptors are suspected to play a role in memory, anxiety, depression, and cognitive deficits associated with long-term hypertension. ^16–18^

Hyperaldosteronism is typically caused by bilateral nodular adrenal disease or hyperplasia, and in 30-40% of cases by unilateral APA. Adrenalectomy is the current standard of care for unilateral disease but is not usually feasible for bilateral disease. For those that can undergo surgery, adrenalectomy is successful in resolving PA, with most complete clinical cure seen in 37% and complete biochemical cure in 94%.^19–22^ However, surgery is not feasible in all patients, including bilateral disease, elderly patients and those with multi-morbidity.

Current standard of care for bilateral disease is medical management with primacy given to mineralocorticoid receptor antagonists (spironolactone or eplerenone).^23^ Spironolactone is not always well tolerated, particularly at higher doses due to off-target effects at the androgen receptor which causes gynecomastia and erectile dysfunction in men and odynomastia in women, In women of reproductive age, spironolactone must also be prescribed with caution as it is not safe during pregnancy.^24,25^ Moreover, in patients with PA for whom MRAs have not adequately de-suppressed renin there is a two-fold increased risk for cardiovascular morbidity when compared to BP-matched primary hypertension.^26^ It is therefore desirable to develop definitive management options for both unilateral and bilateral PA which could potentially spare healthy adrenal tissue and avoid long-term adrenocortical insufficiency.

Targeted thermal ablation represents one such option which could accurately ablate abnormally functioning adrenal adenomas or nodules, while leaving untargeted normally functioning adrenal tissue intact (Quote Padraig Donlon’s Study here IntJTherMed). Currently, thermal ablation of adrenal adenomas is undertaken in selected centres for the management of PA and Cushing Syndrome with the benefits of being minimally invasive. This modality also offers the potential to treat bilateral disease. Percutaneous image-guided approaches to adrenal tumors include radiofrequency ablation, cryoablation, microwave ablation, and chemical ablation. ^27–30^ Microwave ablation utilizes non-ionizing electromagnetic radiation, typically at 915 MHz, 2.45 GHz or 5.8 GHz, to agitate water molecules, producing friction and heat that leads to cellular death and coagulation necrosis. ^31–33^ Microwave ablation, compared to radiofrequency ablation, has a more rapid increase in temperature, can achieve an overall higher local temperature, can target a larger volume, and does not need grounding pads. ^34^ In previous work, our group has developed and tested an *in vivo* porcine model that utilized directed microwave radiation for potential thermal treatment of aldosterone-producing adenomas. ^35^ This work, however, did not include a functioning tumor target, demonstrating the need for a better characterized model of neoplastic PA that is useful for testing thermal therapies.

APA research requires animal models that replicate functional mechanisms of PA to test pre-clinical novel therapeutics. Species differences in both neoplastic conditions and the physiological structure of the adrenal gland are limiting. Humans have three distinct adrenal gland cortical zones: zona glomerulosa, zona fasciculata, and zona reticularis. Aldosterone is produced only in the zona glomerulosa and cortisol in the zona fasciculata due to the exclusive location of their synthesis enzymes, CYP11B2 and CYP11B1, respectively. Mice lack a zona reticularis and thus do not produce adrenal androgens post-puberty. Gene sequencing has shown specific human somatic mutations identified in the APA exome when compared to the germline exome. ^36–38^ While mouse models featuring some genetic alterations could mimic human somatic mutations, the resulting neoplasia may not be similar enough to the human genotype to predict external validity. Low animal-to-human translational success rates can partially be attributed to low predictability of human outcome based on animal physiology, genetics, epigenetics, and molecular biology. ^39,40^ Xenograft models are being implemented into research methods more frequently to increase translational success. These models propose some difficulties, such as management of immunodeficient mice and short life expectancy limiting long-term assessment of therapeutic strategies.

This study characterizes a mouse model that can partially replicate the genetic component of human aldosterone-producing adenomas by utilizing HAC15 cells inoculated into immunodeficient mice. This clonal cell line was derived from a 48-year-old patient with adrenocortical carcinoma and has been shown to be responsive to changes in angiotensin II, potassium, ACTH, and cAMP. HAC15 tumors can secrete mineralocorticoids, glucocorticoids, and adrenal androgens, though this mouse tumor model only produces aldosterone from angiotensin II stimulation. ^41,42^ The HAC15 cell line has also shown expression of the five forms of cytochrome P450 enzymes involved in adrenal steroidogenesis. ^42–45^ This study aims to characterize a PA mouse model useful for assessing biochemical outcomes of thermal therapies.

## 2. Materials and Methods

### 2.1 Materials

The HAC15 human adrenocortical carcinoma cell line was purchased from ATCC (Manassas, VA). General chemicals, Trypan Blue, and DMSO were purchased from Sigma-Aldrich (St. Louis, MO). DMEM/F12 HEPES buffer was manufactured by Life Technologies Corp (Grand Island, NY) and purchased from Fisher Scientific (Waltham, MA). 10% Cosmic Calf Serum was manufactured by Hyclone Laboratories Inc. (Logan, UT) and purchased from Fisher Scientific (Waltham, MA). Phenol red-free Matrigel HC (high concentration) basement membrane matrix, ITS+ premix culture supplement, and phosphate-buffered saline (PBS) were manufactured by Corning Life Sciences (Corning, NY) and purchased from Fisher Scientific (Waltham, MA). 1% penicillin-streptomycin and trypsin-EDTA were purchased from Gibco (Waltham, MA). Heparin was purchased from the Veterinary Health Center at Kansas State University (Manhattan, KS). Angiotensin II >98% purity was purchased from MP Biomedicals. Male athymic nude mice (Nu/Nu) were purchased from Charles River (strain 490) at 4-6 weeks old (Wilmington, MA).

### 2.2 Cell Culture and Preparation

HAC15 cells were stored in a cryopreservation medium consisting of 10% DMSO, 40% complete growth medium (consisting of 1% ITS+ premix, 1% penicillin-streptomycin, 10% cosmic calf serum, and 88% DMEM/F12), and 50% cosmic calf serum in liquid nitrogen vapor. Frozen vials were thawed via a slight agitation in a 37 °C water bath, keeping the cap out of the water. Thawed cells were transferred from the cryovial to a 15 mL centrifuge tube containing 5 mL of culture medium, then the tube was centrifuged at 1,000 rpm (190g) for 3 minutes before the supernatant was removed and the pellet was resuspended with culture medium. Culture medium consisted of 88% DMEM-F12 (Dulbecco’s Modified Eagle Medium), 1% ITS+ (Insulin-Transferrin-Selenium), 1% penicillin-streptomycin, and 10% cosmic calf serum.

Cells were cultured to 80% confluency and split in a 1:2-1:3 density reduction. This is a slow-growing cell line with incomplete adherence. To speed up population growth, all old medium was spun down before disposal with the pellet added back to the flask. To subculture, PBS was used as a plate rinse and trypsin-EDTA was used for lifting cells.

To prepare for mouse injections, cells were lifted with 0.1% trypsin-EDTA and stopped with medium after approximately 5 minutes. The total volume is centrifuged at 1000 rpm (190g) for 3 minutes and the cell pellet was resuspended in ice-cold saline. The cells were stained with trypan blue and counted using a Nexcelom Cellometer on a BXPC-3 cell template. Total injection volume per mouse was 250 µl maximum. Studies that used Matrigel had injection volumes that consisted of ⅔ cell suspension and ⅓ Matrigel HC.

### 2.3 Animal Research and Ethics Statement

All three studies were carried out in strict accordance with the recommendations in the <u>Guide for the Care and Use of Laboratory Animals</u> of the National Academies Press. The protocol was approved by Kansas State University’s Institutional Animal Care and Use Committee (protocol number: 4451).

### 2.4 Tumor Size Experiment

Male, 4 to 6-month-old, athymic nude mice were injected subcutaneously with 250 µl of various cell concentrations +/- Matrigel. Fifteen mice were split into 3 experimental groups. The first group received 6 million HAC15 cells, the second received 10 million HAC15 cells with Matrigel, and the third received 20 million HAC15 cells with Matrigel. Mice were placed in dorsal recumbency under isoflurane anesthesia and HAC15 injections were given subcutaneously into the right hind flank using freezer-chilled 1 ml syringes. Tumor size was measured approximately every other day starting at day 7 post-injection.

### 2.5 Angiotensin-Aldosterone Delay Study

Fourteen male, 4 to 6-month-old, athymic nude mice were injected subcutaneously with 20 million HAC15 cells suspended in Matrigel following protocol from the tumor size experiment. Tumor size was measured every other day. The resulting APAs were grown to at least 5 mm in one dimension at which point each mouse, except 2 control mice which received PBS injections, was held in dorsal recumbency, and given angiotensin II by intraperitoneal injection at a concentration of 1 mg/ml and a dose of 5 µl/g. Following administration of angiotensin II, mice were assigned to a waiting interval of 30 minutes, 1 hour, 2 hours, or 4 hours with 3 mice per interval. At the end of the waiting interval the mice were euthanized via a carbon dioxide chamber and a cardiac puncture was performed with 1 ml heparinized syringes with 25g 5/8” needles. Blood samples were centrifuged at 2,000 x g for 15 minutes. Plasma was removed and sent to Birmingham University, UK for analysis.

### 2.6 Tumor Size Correlation to Aldosterone Production

Eighteen male, 4 to 6-month-old, athymic nude mice were injected with 20 million HAC15 cells suspended in Matrigel following protocol from the tumor size experiment. Tumor size was measured every other day. Three mice were assigned to one of 6 groups. On days 3, 6, 10, 15, 20, or 25 post-HAC15 injection, mice were held in dorsal recumbency and administered angiotensin II by intraperitoneal injection at a concentration of 1 mg/ml and at a dose of 5 µl/g. Two hours post-injection mice were euthanized via a carbon dioxide chamber and a cardiac puncture was performed with 1 ml heparinized syringes. Blood samples were centrifuged at 2,000g for 15 minutes, and plasma removed and frozen. Tumors were excised, measured, flash frozen with liquid nitrogen, and stored at –80°C. Plasma samples and excised tumors were sent to Birmingham University, UK for analysis.

### 2.7 Steroid quantitation by liquid chromatography tandem mass spectrometry

Plasma steroids for the quantitation of aldosterone, progesterone, deoxycorticosterone and corticosterone were extracted using a previously validated method described elsewhere. ^40^ In brief, steroid were extracted from 1-200 μL of plasma after addition of internal standards. Firstly proteins were precipitated with 50 μL of acetonitrile followed by steroid extraction via liquid-liquid extraction with 1 mL tert-methyl butyl ether (MTBE, Acros Organics, Fisher Scientific, UK). The MTBE layer was then removed, transferred into a 96 well plate and evaporated under nitrogen at 30°C. Samples were reconstituted in 125 μL of 50/50 methanol/water and 20 μL was injected into a liquid chromatography tandem mass spectrometer, Waters Acquity with Xevo-XS (Waters Ltd). Steroids were separated on a Phenomenex luna column, C_18_ 2.5 x 50 mm (Phenomenex, UK) at 60°C, using a methanol/ water (0.1% formic acid gradient system) with post-column infusion of 6mM ammonium fluoride.

Steroids were quantified in positive ionization mode using 2 mass transitions; aldosterone 361 > 343 and 361 > 315 (Steraloids, US); progesterone 315 > 97 and 315 > 109 (Sigma Aldrich, UK); deoxycorticosterone 331 > 97 and 331 > 109 (Sigma Aldrich, UK); and corticosterone 347 > 121 and 347 > 97 (Sigma Aldrich, UK). Steroids were quantified relative to their isotopically labelled internal standards aldosterone-d8 (Isosciences, UK), progesterone-d9 (Cambridge Isotope Laboratories, UK), deoxycorticosterone-d8 (Steraloids, US) and corticosterone-d8 (Cambridge Isotope Laboratories, UK) using a calibration range of 0.05 to 250 ng/mL.

### 2.8 RNA Isolation and Gene Expression Analysis on Angiotensin II-Stimulated Mouse Tumors

Mouse tumors, excised at various timepoints and stored at -80°C, were thawed in RNA-later-ICE (Invitrogen™) at -20°C and sectioned into 10 mg wedges, to include cells from the core and periphery of the tumor sample. RNA was isolated from the sections using Qiagen EZ2 Connect (Qiagen) according to the manufacturer’s protocol. RNA integrity and concentration were measured using DeNovix DX-11 Spectrophotometer (DeNovix Inc,) and RNA was stored at - 80°C prior to RT-PCR analysis.

Reverse transcription was performed using Oligo (DT) primers (1µg) and 10nM dNTPs. Samples (1µg) were denatured by incubation at 65°C for 5 min, followed by addition of Superscript III reverse transcriptase (Invitrogen™), 0.1 M 1,4-Dithiothreitol (DTT) and 5X Reaction buffer (Invitrogen™). The reaction mix was incubated at 25°C for 5 min, 50°C for 60 min, and 70°C to denature the enzyme using DNA Engine Dyad thermal cycler (Bio-Rad Laboratories). The synthesized cDNA was stored at -20°C for further use.

Gene expression analysis was performed using GoTaq qPCR SYBR mastermix (Promega), according to manufacturer’s protocol, targeted against CYP11A1, CYP11B1, CYP11B2 and StAR. Housekeeping genes PMM1, HPRT1, RPLP0 were used for normalization. cDNA from HAC15 cells targeting PMM1 was used as inter-assay control. The RT-PCR was performed at 95°C for 10 min followed by 40 cycles of 95°C for 15 secs, 60°C for 1 min using StepOne Plus™ (Applied Biosystems™). The relative quantity (RQ)/ fold change was calculated using the comparative CT method (2-ΔΔCt). ^47^

### 2.9 Treating Tumors with Microwave Ablation

To determine whether this model could be used to evaluate local therapies, twelve mice bearing HAC15 tumors were treated with microwave ablation. Six million HAC15 cells were suspended in 50 μL of PBS and injected subcutaneously into the right hind flank of twelve-week-old male nude mice. Tumor size was measured by caliper every other day starting on day eight post-injection. Tumors were allowed to grow until they reached 5 mm in greatest dimension, at which time mice were assigned to either treatment or control groups (six mice in each group). Mice in the treatment group were anesthetized with isoflurane and the tumor was heated to temperatures of 55 °C or higher with a custom 2.45 GHz microwave antenna using MRI thermometry feedback as previously described. ^35^ Similar to the treatment mice, control group mice were anesthetized with isoflurane and imaged by MRI, but did not receive microwave ablation. Twenty-four hours after treatment or mock-treatment, mice were euthanized using a CO_2_ chamber and tumors, lung, liver, kidney, spleen, small intestine, and brain were collected. Tumors were immediately submerged in a triphenyl tetrazolium chloride (TTC) solution for 1 hour to give immediate feedback as to whether there were functioning (live) mitochondria in the target area – live mitochondria will turn the TTZ stain red, staining live tissue bright red. ^48,49^ Tumors and other tissues were then fixed in 10% buffered neutral formalin for histological analysis.

To determine whether this model could be used to provide biochemical feedback on tumor function after treatment, fourteen mice bearing HAC15 tumors were treated with microwave ablation. Twenty million HAC15 cells were suspended in 167 μL of ice-cold PBS and combined with 83 μL of Matrigel HC for a total injection volume of 250 μL. Injections were given subcutaneously into the right hind flank of twelve-week-old male nude mice. Twenty-five days after injection, mice were split into treatment and control groups (seven mice per group). Treatment mice were anesthetized with isoflurane and had a fiber optic thermal probe inserted into the tumor using a 20g needle; the tumor was then heated using the same microwave ablation antenna as above. Control mice were anesthetized for equivalent time but did not receive microwave ablation treatment. Twenty-four hours after treatment or mock-treatment, mice were injected intraperitoneally with angiotensin II (1 mg/kg in PBS). Two hours after angiotensin II injection, mice were euthanized using a CO_2_ chamber and blood was drawn immediately post-euthanasia by cardiac puncture using heparin-coated syringes. Collected blood was centrifuged at 2000 x g for 15 minutes to separate plasma. Plasma was stored at -20°C for further analysis.

### 2.10 Histology of Tumors and Other Tissues

Formalin-fixed tumors, lung, liver, kidney, spleen, small intestine, and brain were embedded in paraffin, sectioned at 5 µm thickness, placed onto glass slides, and stained with hematoxylin and eosin (H&E) for histologic analysis by light microscopy.

### 2.11 Statistical Methods

All analyses were performed using XLSTAT (2007) Statistical Software for Excel. Differences between the two groups were analyzed using a two-tailed Student’s *t*-test. Depending on whether the assumption of normal distribution and homogeneity of variance was met, an ANOVA or Kruskal-Wallis test was used and succeeded by one of the following post-hoc tests: Tukey’s Honest Significant Difference if parametric or Dunn’s Multiple Comparison if nonparametric. When comparing multiple treatments over time, an ANOVA repeated measures test was utilized.

## 3. Results

### 3.1 Tumor Growth

Three different cell concentrations for inoculation were utilized along with the addition of Matrigel in two of the three groups. Tumor size data (figure 1) demonstrated that both the 10 and 20 million cells plus Matrigel preparations grew significantly faster than 6 million cells without Matrigel and had more homogeneity than the standard six million cells. Statistical analysis using a repeated measures test yielded significant P-values for all time points except 25- and 31-days post-injection. A Tukey’s Honest Significant Difference post-hoc test showed consistent significance between the 20 million cells with Matrigel group and the 6 million cells without Matrigel for every time point. The 10 million cells with Matrigel group had significance when compared to 6 million cells without Matrigel at time points 7-, 11-, 14-, 16-, 18-, and 21-days post-injection. Lastly, the group containing 10 million cells with Matrigel group had significance when compared to the 20 million cells with Matrigel group at time points 11-, 14-, 16-, and 18-days post-injection.

**Fig 1:**
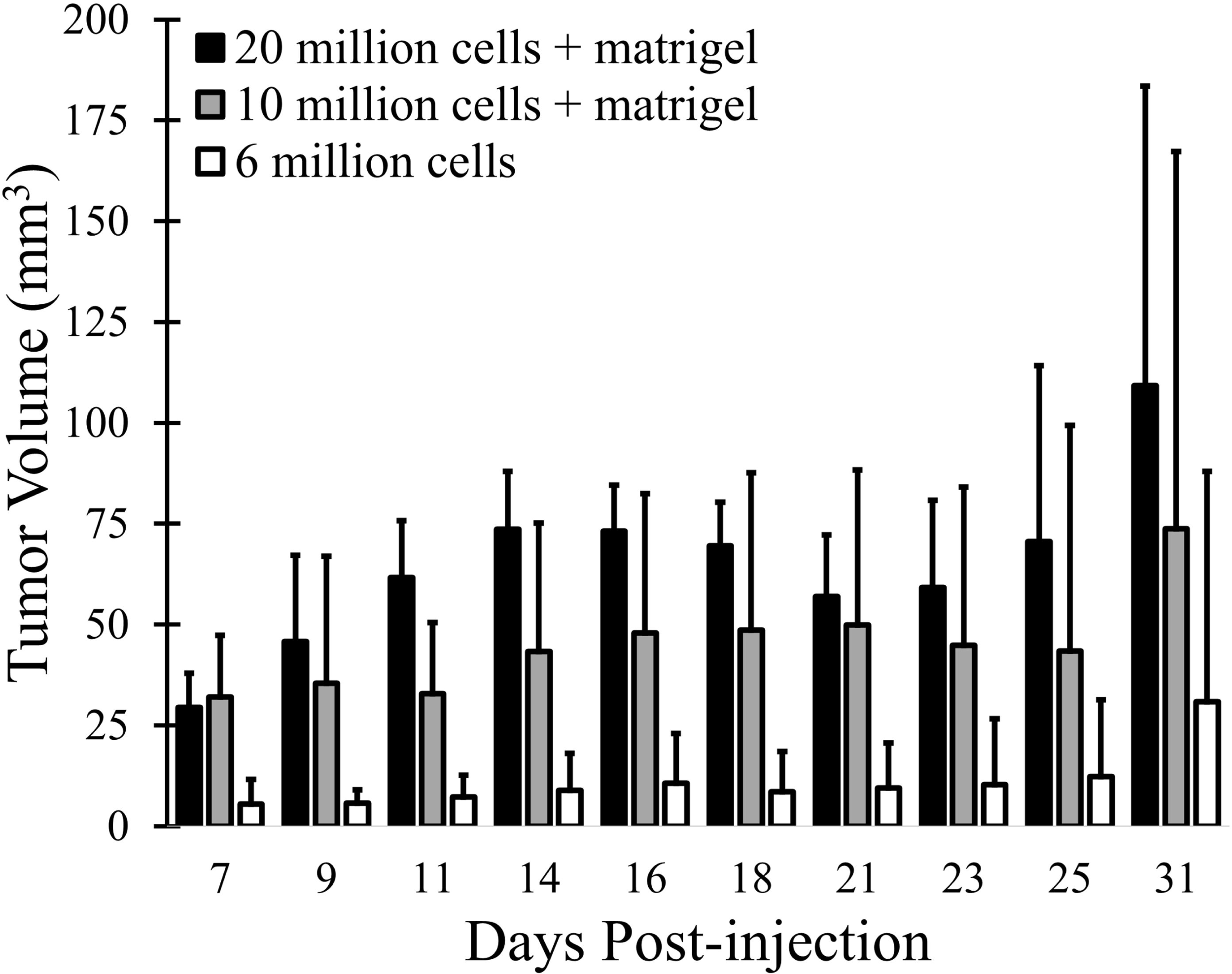
Tumor volume growth post-inoculation. Neoplasms grew in the right hind of flank of immunodeficient mice injected subcutaneously with human HAC15 adrenocortical carcinoma cells. The following inoculation treatment groups were tested: 6 million cells without Matrigel, 10 million cells with Matrigel, and 20 million cells with Matrigel.

Mice with 6 million HAC15 cells injected subcutaneously underwent tumor growth to at least 5 mm in any direction before being euthanized and having the tumor and major organs excised. No evidence of metastasis was identified on histologic evaluation of H&E-stained sections of intestine, brain, lungs, kidneys, spleen, and liver collected from each mouse. Histologically, the subcutaneous tumors were incompletely pseudoencapsulated, multilobular, and formed by densely cellular polygonal cells arranged in cords, nests, and packets supported by a fine fibrovascular stroma. Neoplastic cells had indistinct borders, a moderate amount of eosinophilic to amphophilic, often microvacuolated, cytoplasm, and a round to oval nucleus with finely stippled chromatin and variably sized deeply basophilic nucleoli. Anisocytosis and anisokaryosis were mild to moderate and the mitotic count varied from3 to 12 per 0.255 sq mm field in each tumors. (figure 2).

**Fig 2:**
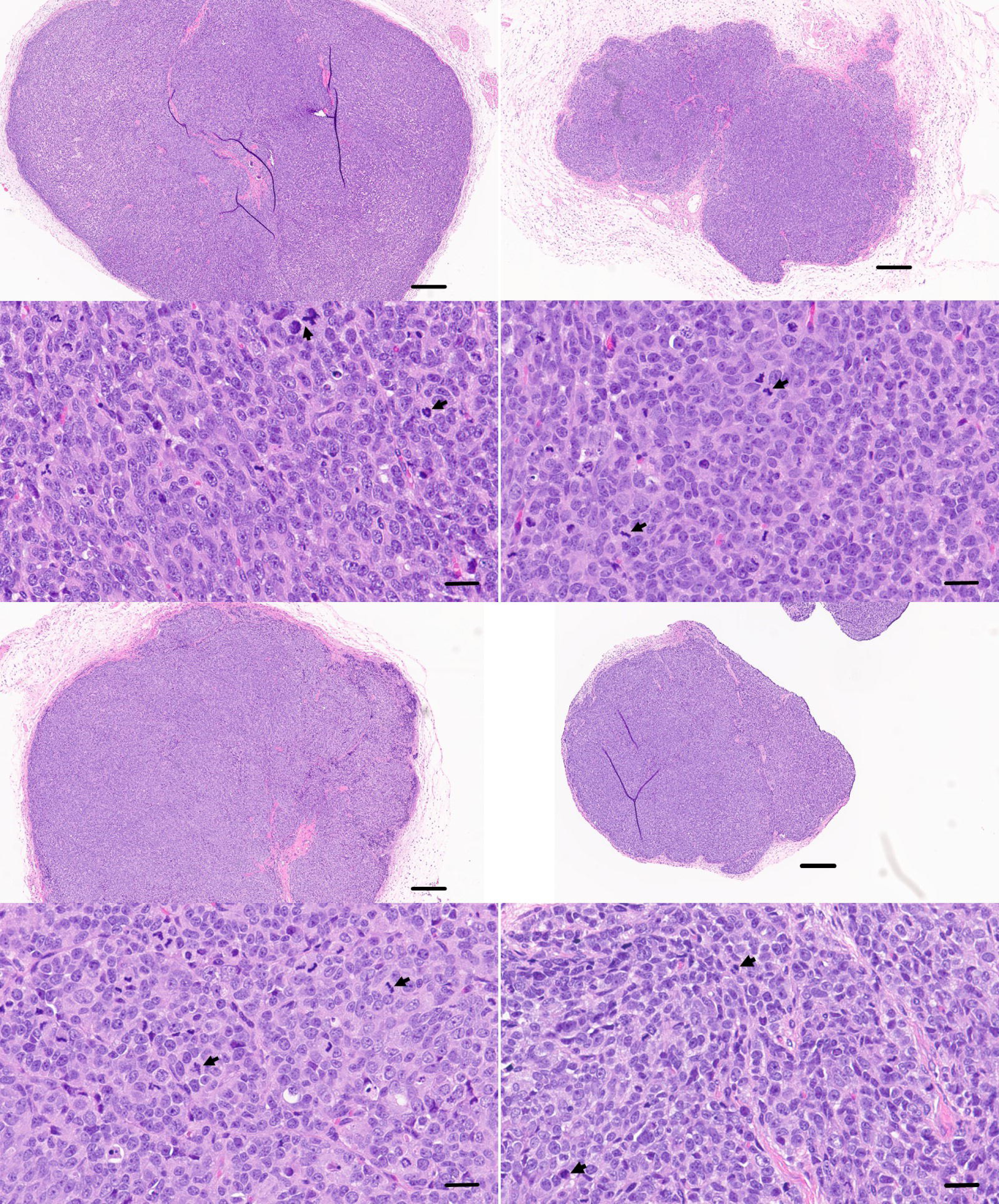
Histology of HAC15 tumor. Tumors grown in the flank of immunodeficient mice injected subcutaneously with human adrenocortical carcinoma (HAC15) cells. H&E stain. Top images:40X magnification, 250 µm scale bars. Bottom images: 400X magnification, 250 µm scale bars; arrows point to mitotic figures.

### 3.2 Response to Angiotensin II

Mice bearing HAC15 tumors and challenged with angiotensin II showed a significant increase relative to baseline (time 0) in aldosterone precursors at 60 or 120 minutes (figure 3a) and a significant increase in plasma aldosterone at 120 minutes (figure 3b). To verify that the increase in steroid concentrations was due to the tumor, mice without tumors were compared to mice with tumors. At 120 minutes post-angiotensin II injections, significant differences were detected between tumor-bearing mice and non-tumor-bearing mice in plasma progesterone (figure 3c), plasma deoxycorticosterone (figure 3d) and plasma aldosterone (figure 3e).

**Fig 3:**
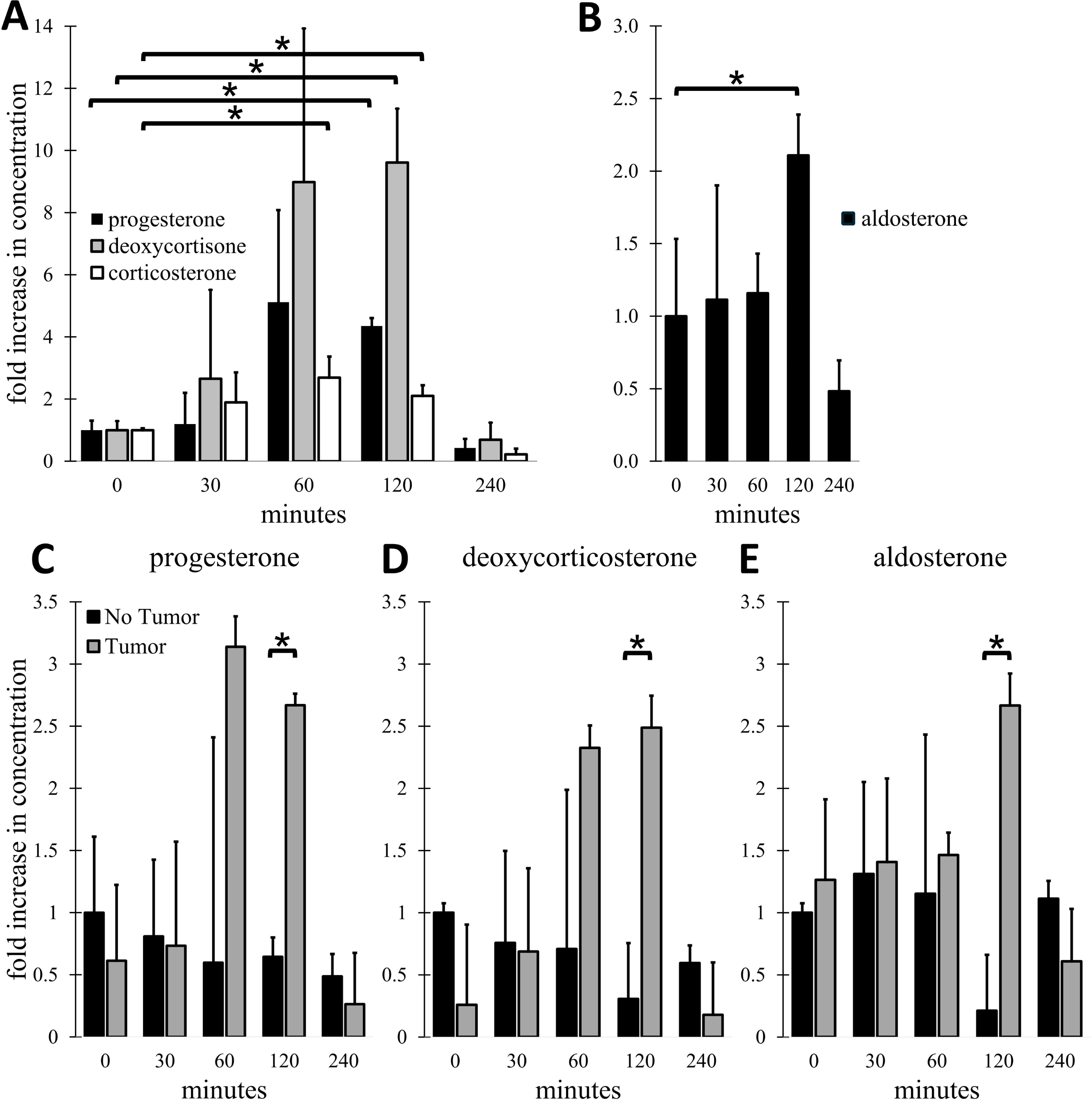
Plasma aldosterone and precursor concentrations post-angiotensin II injections. **(A)** Aldosterone precursors: mice bearing 5 mm HAC15 tumors were given angiotensin II (5μg/g) and then euthanized 30, 60, 90, 120, or 240 minutes afterward to determine at what time point peak precursor concentrations were reached; **(B)** Aldosterone: mice bearing 5 mm HAC15 tumors were given angiotensin II (5μg/g) and then euthanized 30, 60, 90, 120, or 240 minutes afterward to determine at what time point peak aldosterone concentration was reached; **(C-E)** Mice bearing 5 mm HAC15 tumors compared to non-tumor bearing mice; **(C)** progesterone concentrations**; (D)** deoxycorticosterone concentrations**; (E)** aldosterone concentrations. * = significant difference (p<0.05).

### 3.3 Tumor Steroidogenesis

Excised HAC15 tumors showed increased expression of the steroidogenic enzymes relevant for mineralocorticoid and glucocorticoid biosynthesis (CYP11B1, CYP11B2, StAR, CYP11A1) versus controls, although only a few time points were significant (figure 4a-d). Steroidogenic enzyme levels did not correlate well with plasma aldosterone levels measured in the mice (figure 4e), although there was a significant correlation between the change in aldosterone concentration between timepoints to the expression of CYP11B2, with the largest increase in aldosterone concentration matching the peak of measured CYP11B2 expression on day 6 (Pearson Correlation = 0.757, p=0.0407). There was a similar correlation between change in aldosterone concentration to the expression of CYP11A1 (Pearson Correlation = 0.741, p=0.0461). Steroid production correlation to tumor size was also evaluated. No significant correlation between tumor size and aldosterone concentration was noted (figure 4f, Pearson Correlation = -0.1303).

**Fig 4:**
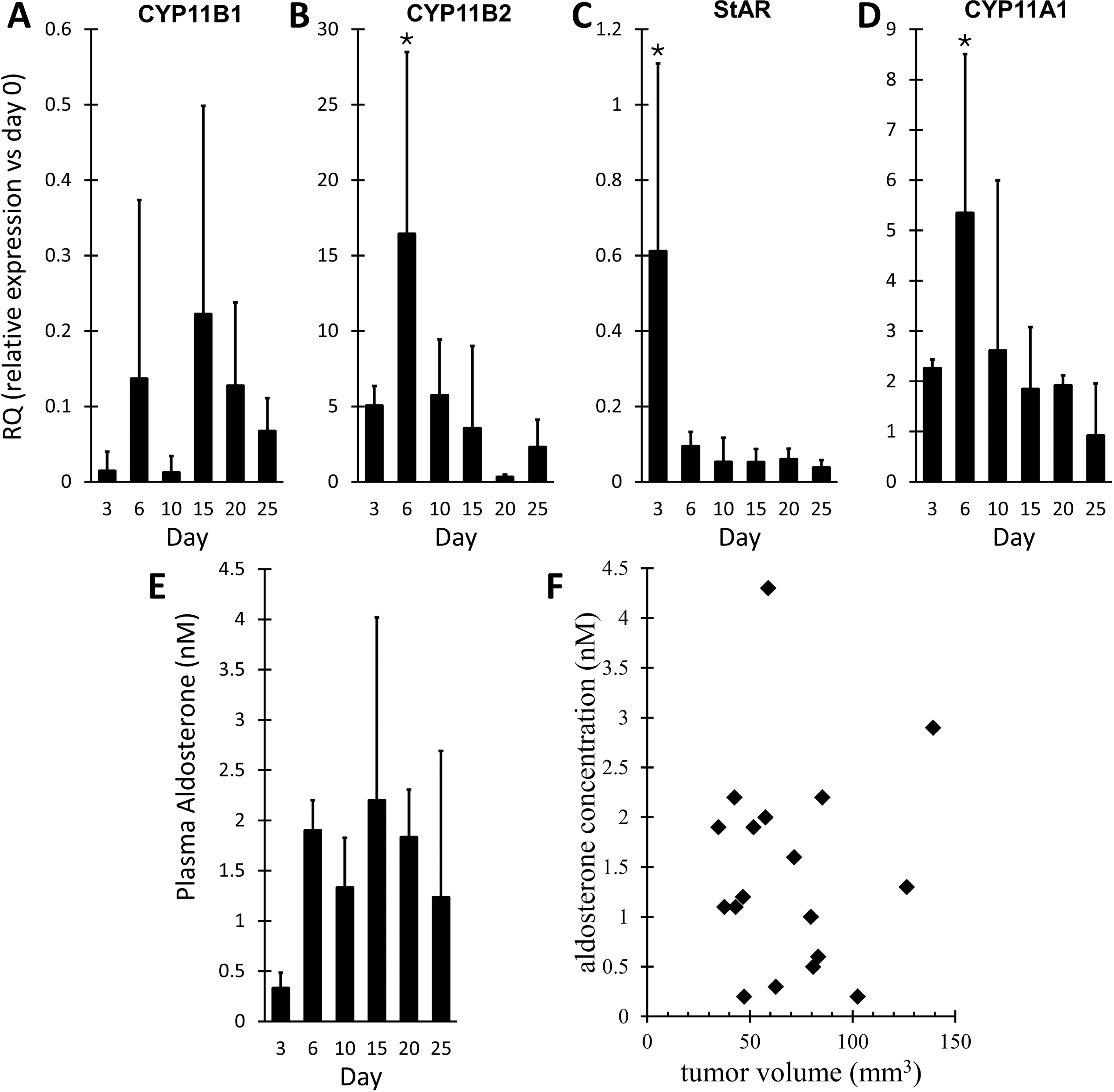
Steroidogenesis. **(A-D)** RT-PCR for steroidogenic enzymes in tumors two hours post-angiotensin II challenge versus days with tumor. **(A)** CYP11B1. **(B)** CYP11B2. **(C)** StAR. **(D)** CYP11A1. **(E)** Plasma aldosterone concentration vs days with tumor. **(F)** Plasma aldosterone concentration versus tumor volume. * = significant difference (p<0.05 vs day 0).

### 3.4 Using the HAC15 Model to Evaluate Treatment with Microwave Ablation

To determine whether this model could be used to evaluate local energy-based ablative therapy, HAC15 tumor-bearing mice were treated with microwave ablation and tumors were collected and analyzed. Tumors that were not ablated stained dark red with TTC, indicating viability (Figure 5b). Most of the ablated tumor tissue did not stain red, indicating loss of tissue viability post-ablation. Small areas of tumors that stained red indicate incomplete tumor ablation due to imperfect targeting of the tumor. Histologically, ablated tumors were less cell dense, with clear space often separating tumor cells (Figure 5a). Ablated tumor cells appeared shrunken and approximately half the diameter of unablated tumor cells. While cell structure was maintained, likely due to thermal fixation, ablated cell cytoplasm was strikingly more eosinophilic and nuclei were more elongate and either uniformly deeply basophilic or exhibited chromatin clumping.

**Fig 5:**
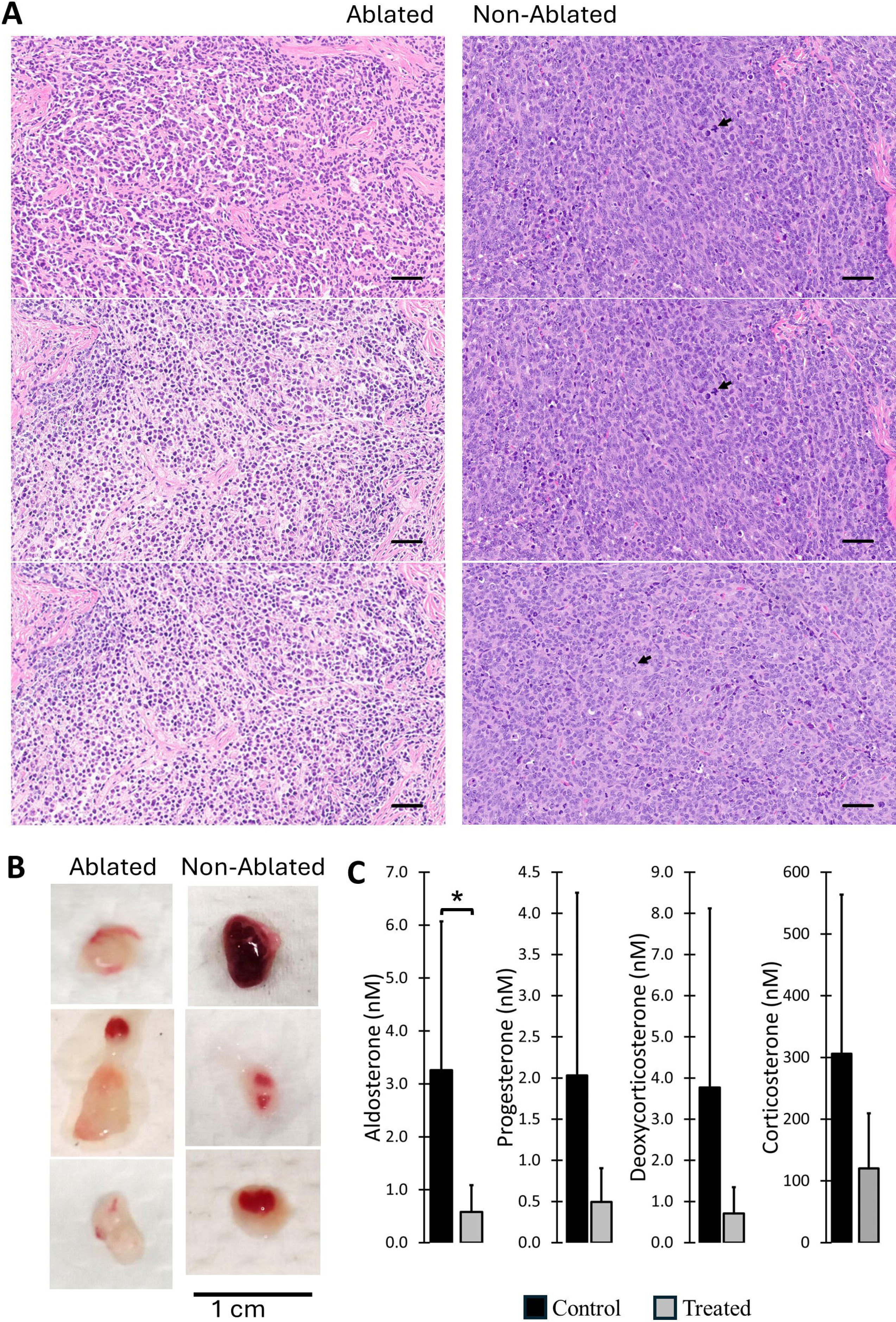
Response to Thermal Ablation Treatment. **(A)** Representative H&E-stained sections of tumors harvested 24 hours post-ablation (left column) compared to non-ablated tumors (right column). Each image taken at 200X; scale bars: 50 µm; arrows point to mitotic figures.. **(B)** Representative images of viability stain 24 hours post-ablation (red = metabolically active tissue). **(C)** Plasma aldosterone and precursors 24 hours post-ablation and two hours post-angiotensin II challenge in treated and untreated (control) mice. * = significant difference (p<0.05).

An effective APA model should allow correlation of treatment status and steroid production. To demonstrate this in the HAC15 model, HAC15 tumor-bearing mice were treated with microwave ablation and twenty-four hours later the steroidogenesis of the tumor was analyzed by challenge with angiotensin II. Plasma aldosterone levels in treated mice were significantly lower than in untreated mice (figure 5c). Similar reductions in other steroid precursors (progesterone, deoxycorticosterone, corticosterone) also occurred but lacked statistical significance due to large standard deviations.

## 4. Discussion

This study demonstrates the development and characterization of a mouse model of human primary neoplastic aldosteronism that is useful for testing ablative therapies for aldosterone producing adenomas (APAs). An optimal model to test thermal therapies,must replicate basic features of human pathophysiology^50^ and be able to characterize the change in blood biochemistry before and after treatment. Multiple mouse models of PA currently exist, although the utility of current mouse models has been questioned due to significant differences between the pathophysiology of mouse tumors and human tumors. ^50^ A xenograft mouse model of human metastatic adrenocortical carcinoma has been previously developed and characterized using the human H295R cell line. ^51,52^ The HAC15 cell line is a derivative of the H295R cell line that was selected for responsiveness to both angiotensin II and ACTH. ^42^ When compared to the H295R line, HAC15 demonstrates a broader spectrum of steroidogenesis, ^41^ sensitivity to ACTH and angiotensin II, ^42^ and a more stable steroidogenic phenotype, ^41^ suggesting that the HAC15 line could be useful for developing a xenograft mouse model of PA. ^53^

True models of APA should be non-malignant – metastatic aldosteronism would be classified as aldosterone-secreting adrenocortical carcinomas (ACC). The parent H295R line was derived from an ACC patient, but previous subcutaneous models using H295R cells did not demonstrate malignant potential. ^42^ Similarly, during this study, the HAC15 model did not metastasize, suggesting a model representative of APA when used within the time course described herein.

One surprising result from this study was the apparent disconnect between aldosterone production and tumor size in the model (fig. 4f). This may not be a weakness of the model, though, as currently the literature is significantly split on whether size and aldosterone production correlate in APAs. Several studies have suggested that tumor size correlates to aldosterone production and symptoms, ^54,55^ while other studies suggest that there is no correlation between size, aldosterone production, and symptoms in human APA patients. ^56–58^

The HAC15 model also proved to be useful for analyzing a local microwave ablation treatment. Physical readouts, such as the viability stain and H&E characterization of the tumors, demonstrated the utility of the ablation treatment. This was matched by the biochemical characterization of the treatment, which demonstrated a significant reduction in plasma aldosterone post-treatment. Since this model replicates several features of human tumors and has readouts that can be used to analyze the utility of local treatments, it can be a useful model of aldosterone-producing adenomas.

In conclusion, the model developed here should be useful for characterizing the effects of ablative thermal treatments on aldosterone producing adenomas and may potentially be useful for analyzing other treatments as well.

## References

1. Hundemer GL, Vaidya A. Primary Aldosteronism Diagnosis and Management: A Clinical Approach. Endocrinol Metab Clin North Am. Dec 2019;48(4):681–700. doi:10.1016/j.ecl.2019.08.002

2. Wang JJ, Peng KY, Wu VC, Tseng FY, Wu KD. CTNNB1 Mutation in Aldosterone Producing Adenoma. Endocrinol Metab (Seoul). Sep 2017;32(3):332–338. doi:10.3803/EnM.2017.32.3.332

3. Wagner CA. Effect of mineralocorticoids on acid-base balance. Nephron Physiol. 2014;128(1-2):26–34. doi:10.1159/000368266

4. Amar L, Plouin PF, Steichen O. Aldosterone-producing adenoma and other surgically correctable forms of primary aldosteronism. Orphanet J Rare Dis. May 19 2010;5:9. doi:10.1186/1750-1172-5-9

5. White PC. Aldosterone: direct effects on and production by the heart. J Clin Endocrinol Metab. Jun 2003;88(6):2376–83. doi:10.1210/jc.2003-030373

6. Catena C, Colussi G, Nadalini E, et al. Cardiovascular outcomes in patients with primary aldosteronism after treatment. Arch Intern Med. Jan 14 2008;168(1):80–5. doi:10.1001/archinternmed.2007.33

7. Monticone S, Burrello J, Tizzani D, et al. Prevalence and Clinical Manifestations of Primary Aldosteronism Encountered in Primary Care Practice. J Am Coll Cardiol. Apr 11 2017;69(14):1811–1820. doi:10.1016/j.jacc.2017.01.052

8. Chen W, Li F, He C, Zhu Y, Tan W. Elevated prevalence of abnormal glucose metabolism in patients with primary aldosteronism: a meta-analysis. Ir J Med Sci. Jun 2014;183(2):283–91. doi:10.1007/s11845-013-1007-x

9. Hanslik G, Wallaschofski H, Dietz A, et al. Increased prevalence of diabetes mellitus and the metabolic syndrome in patients with primary aldosteronism of the German Conn’s Registry. Eur J Endocrinol. Nov 2015;173(5):665–75. doi:10.1530/eje-15-0450

10. Arlt W, Lang K, Sitch AJ, et al. Steroid metabolome analysis reveals prevalent glucocorticoid excess in primary aldosteronism. JCI Insight. Apr 20 2017;2(8)doi:10.1172/jci.insight.93136

11. Mansour N, Bruedgam D, Dischinger U, et al. Effect of mild cortisol cosecretion on body composition and metabolic parameters in patients with primary hyperaldosteronism. Clin Endocrinol (Oxf). Mar 2024;100(3):212–220. doi:10.1111/cen.15013

12. Wu VC, Chan CK, Wu WC, et al. New-onset diabetes mellitus risk associated with concurrent autonomous cortisol secretion in patients with primary aldosteronism. Hypertens Res. Feb 2023;46(2):445–455. doi:10.1038/s41440-022-01086-w

13. Spyroglou A, Handgriff L, Müller L, et al. The metabolic phenotype of patients with primary aldosteronism: impact of subtype and sex - a multicenter-study of 3566 Caucasian and Asian subjects. Eur J Endocrinol. Sep 1 2022;187(3):361–372. doi:10.1530/eje-22-0040

14. Katabami T, Matsuba R, Kobayashi H, et al. Primary aldosteronism with mild autonomous cortisol secretion increases renal complication risk. Eur J Endocrinol. Apr 25 2022;186(6):645–655. doi:10.1530/eje-21-1131

15. Gerards J, Heinrich DA, Adolf C, et al. Impaired Glucose Metabolism in Primary Aldosteronism Is Associated With Cortisol Cosecretion. J Clin Endocrinol Metab. Aug 1 2019;104(8):3192–3202. doi:10.1210/jc.2019-00299

16. Engler L, Adolf C, Heinrich DA, et al. Effects of chronically high levels of aldosterone on different cognitive dimensions: an investigation in patients with primary aldosteronism. Endocr Connect. Apr 2019;8(4):407–415. doi:10.1530/ec-19-0043

17. Brinks V, van der Mark MH, de Kloet ER, Oitzl MS. Differential MR/GR activation in mice results in emotional states beneficial or impairing for cognition. Neural Plast. 2007;2007:90163. doi:10.1155/2007/90163

18. Hlavacova N, Wes PD, Ondrejcakova M, et al. Subchronic treatment with aldosterone induces depression-like behaviours and gene expression changes relevant to major depressive disorder. Int J Neuropsychopharmacol. Mar 2012;15(2):247–65. doi:10.1017/s1461145711000368

19. Iacobone M, Citton M, Viel G, Rossi GP, Nitti D. Approach to the surgical management of primary aldosteronism. Gland Surg. Feb 2015;4(1):69–81. doi:10.3978/j.issn.2227-684X.2015.01.05

20. Citton M, Viel G, Rossi GP, Mantero F, Nitti D, Iacobone M. Outcome of surgical treatment of primary aldosteronism. Langenbecks Arch Surg. Apr 2015;400(3):325–31. doi:10.1007/s00423-014-1269-4

21. Steichen O, Zinzindohoué F, Plouin PF, Amar L. Outcomes of adrenalectomy in patients with unilateral primary aldosteronism: a review. Horm Metab Res. Mar 2012;44(3):221–7. doi:10.1055/s-0031-1299681

22. Lumachi F, Ermani M, Basso SM, Armanini D, Iacobone M, Favia G. Long-term results of adrenalectomy in patients with aldosterone-producing adenomas: multivariate analysis of factors affecting unresolved hypertension and review of the literature. Am Surg. Oct 2005;71(10):864–9.

23. Connell JM, MacKenzie SM, Freel EM, Fraser R, Davies E. A lifetime of aldosterone excess: long-term consequences of altered regulation of aldosterone production for cardiovascular function. Endocr Rev. Apr 2008;29(2):133–54. doi:10.1210/er.2007-0030

24. Stripp B, Taylor AA, Bartter FC, et al. Effect of spironolactone on sex hormones in man. J Clin Endocrinol Metab. Oct 1975;41(4):777–81. doi:10.1210/jcem-41-4-777

25. Kim GK, Del Rosso JQ. Oral Spironolactone in Post-teenage Female Patients with Acne Vulgaris: Practical Considerations for the Clinician Based on Current Data and Clinical Experience. J Clin Aesthet Dermatol. Mar 2012;5(3):37–50.

26. Hundemer GL, Curhan GC, Yozamp N, Wang M, Vaidya A. Cardiometabolic outcomes and mortality in medically treated primary aldosteronism: a retrospective cohort study. Lancet Diabetes Endocrinol. Jan 2018;6(1):51–59. doi:10.1016/s2213-8587(17)30367-4

27. Munver R, Del Pizzo JJ, Sosa RE. Adrenal-preserving minimally invasive surgery: the role of laparoscopic partial adrenalectomy, cryosurgery, and radiofrequency ablation of the adrenal gland. Curr Urol Rep. Feb 2003;4(1):87–92. doi:10.1007/s11934-003-0065-4

28. Xiao YY, Tian JL, Li JK, Yang L, Zhang JS. CT-guided percutaneous chemical ablation of adrenal neoplasms. AJR Am J Roentgenol. Jan 2008;190(1):105–10. doi:10.2214/ajr.07.2145

29. Maki DD, Haskal ZJ, Matthies A, et al. Percutaneous ethanol ablation of an adrenal tumor. AJR Am J Roentgenol. Apr 2000;174(4):1031–2. doi:10.2214/ajr.174.4.1741031

30. Venkatesan AM, Locklin J, Dupuy DE, Wood BJ. Percutaneous ablation of adrenal tumors. Tech Vasc Interv Radiol. Jun 2010;13(2):89–99. doi:10.1053/j.tvir.2010.02.004

31. Uppot RN, Gervais DA. Imaging-guided adrenal tumor ablation. AJR Am J Roentgenol. Jun 2013;200(6):1226–33. doi:10.2214/ajr.12.10328

32. Simon CJ, Dupuy DE, Mayo-Smith WW. Microwave ablation: principles and applications. Radiographics. Oct 2005;25 Suppl 1:S69–83. doi:10.1148/rg.25si055501

33. Ethier MD, Beland MD, Mayo-Smith W. Image-guided ablation of adrenal tumors. Tech Vasc Interv Radiol. Dec 2013;16(4):262–8. doi:10.1053/j.tvir.2013.08.008

34. Yamakado K. Image-guided ablation of adrenal lesions. Semin Intervent Radiol. Jun 2014;31(2):149–56. doi:10.1055/s-0034-1373797

35. Sebek J, Shrestha TB, Basel MT, et al. System for delivering microwave ablation to subcutaneous tumors in small-animals under high-field MRI thermometry guidance. Int J Hyperthermia. 2022;39(1):584–594. doi:10.1080/02656736.2022.2061727

36. Oki K, Plonczynski MW, Luis Lam M, Gomez-Sanchez EP, Gomez-Sanchez CE. Potassium channel mutant KCNJ5 T158A expression in HAC-15 cells increases aldosterone synthesis. Endocrinology. Apr 2012;153(4):1774–82. doi:10.1210/en.2011-1733

37. Spät A, Hunyady L. Control of aldosterone secretion: a model for convergence in cellular signaling pathways. Physiol Rev. Apr 2004;84(2):489–539. doi:10.1152/physrev.00030.2003

38. Choi M, Scholl UI, Yue P, et al. K+ channel mutations in adrenal aldosterone-producing adenomas and hereditary hypertension. Science. Feb 11 2011;331(6018):768–72. doi:10.1126/science.1198785

39. Pound P, Ritskes-Hoitinga M. Is it possible to overcome issues of external validity in preclinical animal research? Why most animal models are bound to fail. J Transl Med. Nov 7 2018;16(1):304. doi:10.1186/s12967-018-1678-1

40. Leenaars CHC, Kouwenaar C, Stafleu FR, et al. Animal to human translation: a systematic scoping review of reported concordance rates. J Transl Med. Jul 15 2019;17(1):223. doi:10.1186/s12967-019-1976-2

41. Wang T, Rainey WE. Human adrenocortical carcinoma cell lines. Molecular and cellular endocrinology. Mar 31 2012;351(1):58–65. doi:10.1016/j.mce.2011.08.041

42. Parmar J, Key RE, Rainey WE. Development of an adrenocorticotropin-responsive human adrenocortical carcinoma cell line. J Clin Endocrinol Metab. Nov 2008;93(11):4542–6. doi:10.1210/jc.2008-0903

43. Denner K, Rainey WE, Pezzi V, Bird IM, Bernhardt R, Mathis JM. Differential regulation of 11 beta-hydroxylase and aldosterone synthase in human adrenocortical H295R cells. Molecular and cellular endocrinology. Jul 23 1996;121(1):87–91. doi:10.1016/0303-7207(96)03853-1

44. Bird IM, Pasquarette MM, Rainey WE, Mason JI. Differential control of 17 alpha-hydroxylase and 3 beta-hydroxysteroid dehydrogenase expression in human adrenocortical H295R cells. J Clin Endocrinol Metab. Jun 1996;81(6):2171–8. doi:10.1210/jcem.81.6.8964847

45. Samandari E, Kempná P, Nuoffer JM, Hofer G, Mullis PE, Flück CE. Human adrenal corticocarcinoma NCI-H295R cells produce more androgens than NCI-H295A cells and differ in 3beta-hydroxysteroid dehydrogenase type 2 and 17,20 lyase activities. J Endocrinol. Dec 2007;195(3):459–72. doi:10.1677/joe-07-0166

46. Schiffer L, Shaheen F, Gilligan LC, et al. Multi-steroid profiling by UHPLC-MS/MS with post-column infusion of ammonium fluoride. J Chromatogr B Analyt Technol Biomed Life Sci. Oct 15 2022;1209:123413. doi:10.1016/j.jchromb.2022.123413

47. Livak KJ, Schmittgen TD. Analysis of relative gene expression data using real-time quantitative PCR and the 2(-Delta Delta C(T)) Method. Methods. Dec 2001;25(4):402–8. doi:10.1006/meth.2001.1262

48. Durick NA, Laeseke PF, Broderick LS, et al. Microwave ablation with triaxial antennas tuned for lung: results in an in vivo porcine model. Radiology. Apr 2008;247(1):80–7. doi:10.1148/radiol.2471062123

49. Shafirstein G, Hennings L, Kaufmann Y, et al. Conductive interstitial thermal therapy (CITT) device evaluation in VX2 rabbit model. Technol Cancer Res Treat. Jun 2007;6(3):235–46. doi:10.1177/153303460700600311

50. Aragao-Santiago L, Gomez-Sanchez CE, Mulatero P, Spyroglou A, Reincke M, Williams TA. Mouse Models of Primary Aldosteronism: From Physiology to Pathophysiology. Endocrinology. Dec 1 2017;158(12):4129–4138. doi:10.1210/en.2017-00637

51. Logié A, Boudou P, Boccon-Gibod L, et al. Establishment and characterization of a human adrenocortical carcinoma xenograft model. Endocrinology. Sep 2000;141(9):3165–71. doi:10.1210/endo.141.9.7668

52. Morin A, Ruggiero C, Robidel E, et al. Establishment of a mouse xenograft model of metastatic adrenocortical carcinoma. Oncotarget. Aug 1 2017;8(31):51050–51057. doi:10.18632/oncotarget.16909

53. Wang T, Rowland JG, Parmar J, Nesterova M, Seki T, Rainey WE. Comparison of aldosterone production among human adrenocortical cell lines. Horm Metab Res. Mar 2012;44(3):245–50. doi:10.1055/s-0031-1298019

54. Nakai K, Manaka K, Sato J, et al. Aldosterone-Producing Adenomas of Increased Size Are Associated With Higher Steroidogenic Activity. J Clin Endocrinol Metab. Nov 23 2022;107(11):3045–3054. doi:10.1210/clinem/dgac530

55. Omura M, Sasano H, Saito J, Yamaguchi K, Kakuta Y, Nishikawa T. Clinical characteristics of aldosterone-producing microadenoma, macroadenoma, and idiopathic hyperaldosteronism in 93 patients with primary aldosteronism. Hypertens Res. Nov 2006;29(11):883–9. doi:10.1291/hypres.29.883

56. Born-Frontsberg E, Reincke M, Beuschlein F, Quinkler M. Tumor size of Conn’s adenoma and comorbidities. Horm Metab Res. Oct 2009;41(10):785–8. doi:10.1055/s-0029-1224200

57. Giacchetti G, Ronconi V, Rilli S, Guerrieri M, Turchi F, Boscaro M. Small tumor size as favorable prognostic factor after adrenalectomy in Conn’s adenoma. Eur J Endocrinol. Apr 2009;160(4):639–46. doi:10.1530/eje-08-0902

58. Nanba K, Tsuiki M, Sawai K, et al. Histopathological diagnosis of primary aldosteronism using CYP11B2 immunohistochemistry. J Clin Endocrinol Metab. Apr 2013;98(4):1567–74. doi:10.1210/jc.2012-3726

